# The statistics of natural shapes predict high-level aftereffects in human vision

**DOI:** 10.1101/2023.01.02.522484

**Authors:** Yaniv Morgenstern, Katherine R. Storrs, Filipp Schmidt, Frieder Hartmann, Henning Tiedemann, Johan Wagemans, Roland W. Fleming

## Abstract

Shape perception is essential for numerous everyday behaviors from object recognition to grasping and handling objects. Yet how the brain encodes shape remains poorly understood. Here, we probed shape representations using visual aftereffects—perceptual distortions that occur following extended exposure to a stimulus—to resolve a long-standing debate about shape encoding. We implemented contrasting low-level and high-level computational models of neural adaptation, which made precise and distinct predictions about the illusory shape distortions the observers experience following adaptation. Directly pitting the predictions of the two models against one another revealed that the perceptual distortions are driven by high-level shape attributes derived from the statistics of natural shapes. Our findings suggest that the diverse shape attributes thought to underlie shape encoding (e.g., curvature distributions, ‘skeletons’, aspect ratio) are the result of a visual system that learns to encode natural shape geometries based on observing many objects.

---

Shape perception is critical for many evolutionarily important tasks, like object recognition (Biederman, 1987; Marr and Nishihara, 1978; Pentland, 1986; Landau, Smith, and Jones, 1998), and in inferring an object’s material properties (Pinna and Deiana, 2015; Paulun, Kawabe, Nishida, and Fleming, 2015; Paulun, Schmidt, van Assen, and Fleming, 2017), causal history (Leyton, 1992; Spröte, Schmidt, and Fleming; 2016; Schmidt, Phillips and Fleming, 2019; Fleming and Schmidt, 2019), or where and how to grasp it (Eloka and Franz, 2011; Kleinholdermann, Franz, and Gegenfurtner, 2013; Klein, Maiello, Paulun, and Fleming, 2020; Culpers, Brenner, and Smeets, 2006). Yet different shapes vary enormously, and there are potentially limitless ways of describing the differences. It remains unclear how shape is represented in the human visual system.

One well-known method for examining shape coding is with adaptation paradigms that produce shape aftereffects. In such experiments, prolonged viewing of one shape (‘adaptor’) makes a subsequently presented shape (‘test’) appear distorted (Regan and Hamstra, 1992; Rivest, Kim, and Sharpe, 2004; Suzuki and Cavanagh, 1998; Suzuki, 2001, 2003, 2005; Fleming, Holtmann-Rice, and Bülthoff, 2011; Storrs and Arnold, 2013, 2017; Bowden et al., 2019). A given test shape can take on multiple different post-adaptation appearances, depending on which shapes are used as adaptors (e.g., Regan and Hamstra, 1992; Suzuki, 2001, 2005; Storrs and Arnold, 2017; Bowden et al., 2019). For example, in **Figure 1A**, we find that after viewing the magenta-labelled shape, participants experienced the central, white-labelled test stimulus to appear significantly more like the green-labelled shape than it did previously. Yet, the same test stimulus appears more like the magenta shape after prolonged viewing of the green shape. Thus, adaptation can make identical stimuli appear different from one another. Conversely, different test shapes can be made to appear more similar when viewed after appropriately chosen adaptors (**Figure 1B**). Such effects—whereby a given stimulus’ appearance can be ‘pushed’ in different directions—suggest an underlying multidimensional ‘shape space’ encoding multiple factors known to be important in shape perception (Op de Beeck, Wagemans, and Vogels, 2001), such as taper (Suzuki and Cavanagh, 1998), curvature (Suzuki, 2001), aspect ratio (Regan and Hamstra, 1992), and spikiness (Haung, 2020; Bao et al., 2020). Probing shape encoding through adaptation experiments opens the possibility of uncovering the feature dimensions underlying such a shape-space, and thus the rationale that the visual system uses to represent shape.

**Figure 1.**
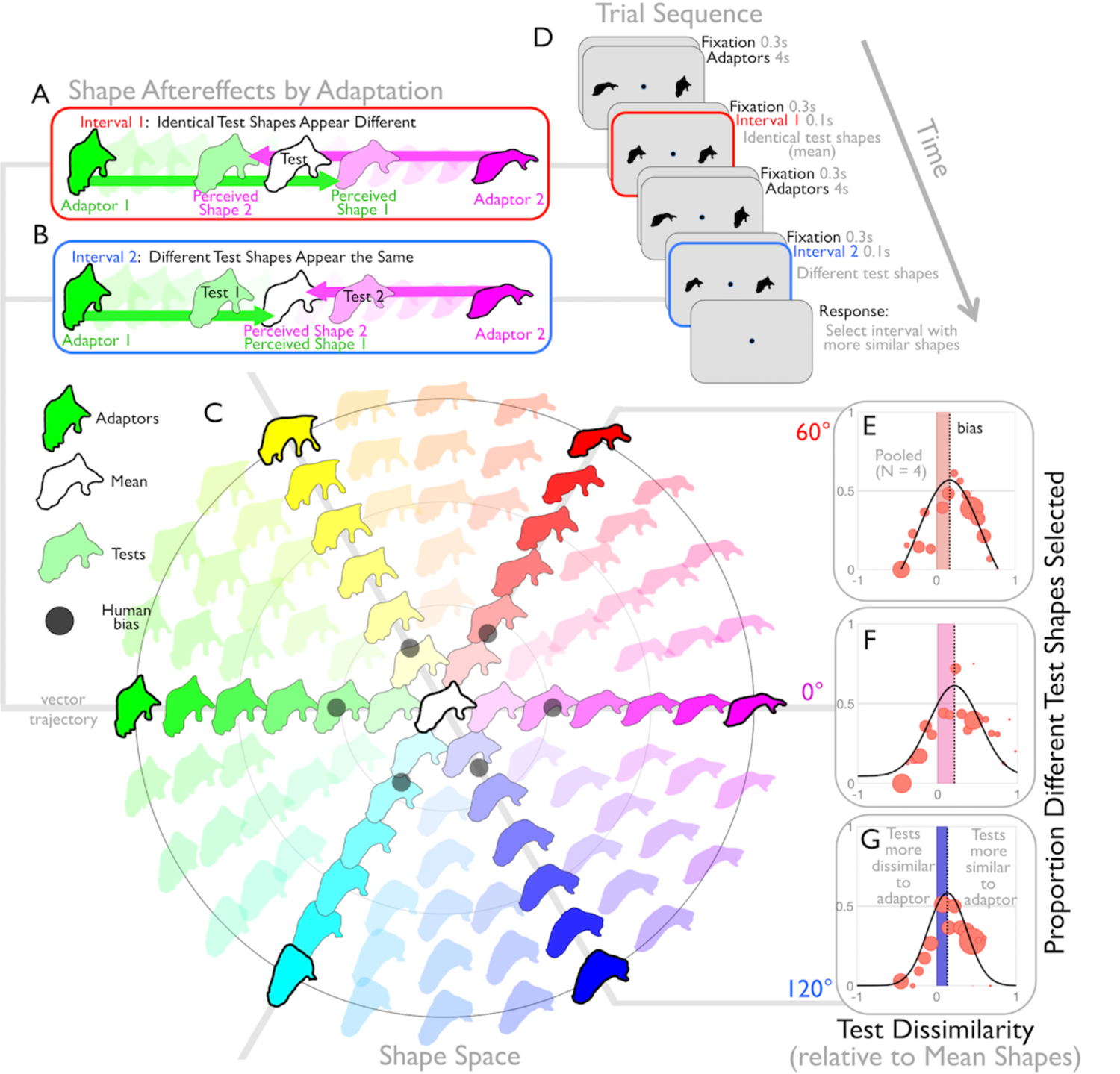
Adaptation along vector trajectories in a statistical shape space yields perceptual aftereffects. Different adapting shapes can make (**A**) an identical test shape appear different, and (**B**) different test shapes appear the same. (**C**) To predict these shifts, we explored adaptation using naturalistic GAN shapes projected onto a 2D slice in a statistical shape space, *ShapeComp,* which represents shape using statistical shape descriptors to summarize the way natural shapes vary (Morgenstern et al., 2021). Shapes are arranged by their ShapeComp coordinates relative to the central *mean shape* (white), with colour saturation indicating dissimilarity from the mean shape and hue reflecting the angular component of the deviation from the mean shape. In separate sessions, the adaptation experiment presented stimuli that passed through single vector trajectories in shape space (at 0°, 60°, or 120°), using adaptors from both ends of the outer ring, and test shapes falling between them. If shape space can predict aftereffects, then shape aftereffects would manifest as a systemic shift away from adaptors on either side of the vector trajectory: an adaptor perceptually distorts the following test shape to appear more like a shape further down the vector trajectory making the mean appear like a shape further away from the adaptor (**A**), and different test shapes appears more like the mean shape (**B**). Thus, by utilizing both adaptors in a 2IFC paradigm (**D**), we hypothesized that we can shift two different test shapes to appear more similar to each other after adaptation than two identical mean shapes (See Movie S1 as an example). We confirmed this by measuring the proportion of (different) test shapes selected over (identical) mean shapes, as a function of test dissimilarity from the mean shape. In the 0°, 60°, and 120° (**E,F,G**) conditions, we find that after the adaptor interval there exists a range of test shapes that observers prefer as more similar than identical mean shapes (black curve shifted to the right; curves are maximum-likelihood fits of a double-sigmoid function through the red data points, see **Materials and methods** for details). The shaded region shows that magnitude of the maximum aftereffect relative to a no adaptation prediction (i.e., zero test dissimilarity), where the varying colours indicate the different vector trajectories. We call the test dissimilarity that leads to the highest probability of selecting different test shapes over identical mean shapes (i.e., peak of the black curve) the ‘shape bias’, which is also shown for each vector trajectory as achromatic dots in (**C**).

A successful model of shape perception should be able to predict how appearance changes after adapting to arbitrary shapes. So-called ‘low-level’ accounts of shape adaptation are based on the early stages of visual processing, where clear links have been found between neural response characteristics and perceptual aftereffects (Hubel and Wiesel 1968; Gibson and Radner, 1937, Blakemore and Sutton, 1969; Mitchell and Muir, 1976, Magnussen and Kurtenbach, 1980, Wilson, McFarlane, and Phillips, 1982; McGraw, Whitaker, Skillen, and Chung, 2002; Bowden et al., 2019). For example, adaptation in populations of orientation-selective cells can explain the well-known ‘tilt aftereffect’ (TAE) in which the orientation of elongated test stimuli (lines, gratings, etc.) appears to shift away from the orientation of an adaptor (Schwartz, Hsu, & Dayan, 2007; Schwartz, Sejnowski, & Dayan, 2009; Clifford, 2014; Dickinson, Harman, Tan, Almeida, & Badcock, 2012; Gibson & Radner,1937; Mitchell & Muir, 1976; Clifford, Wenderoth, & Spehar,2000). By extension, if the perceived orientation of every point on an object’s boundary is distorted to different extents through TAEs induced by the nearby locations on an adaptor stimulus, then the result would be that the entire object appears to have changed shape. Thus, local, low-level processes could in principle explain some or all of the perceived shape distortions.

An alternative account of shape aftereffects is based on later stages of visual processing, where it is thought that higher-level, more global aspects of shape are encoded (Brincat & Connor, 2004; Grill-Spector et al., 2001; Kayaert et al., 2005; Kourtzi & Kanwisher, 2000; Malach et al., 1995). For example, if an adaptor stimulus were thinner and more angular than the test stimulus, neural populations in area IT encoding the ‘thinness’ and ‘angularity’ of the stimuli could adapt, causing the test stimulus to appear more bulbous and curvaceous in comparison. Here, however, the link between aftereffects and neurophysiology is less clear: some argue for figural aftereffects modulated in a high-level shape space (e.g., shape specific dimensions, Suzuki, 2001, 2005; Suzuki & Cavanagh, 1998; faces, Leopold et al., 2001; Leopold & Bondar, 2005; Rhodes & Jeffery, 2006; Rhodes, Jeffery, Clifford, & Leopold, 2007; Webster & MacLeod, 2011; body, Bratch et al., 2021), whereas others have maintained that adaptation aftereffects can be fully explained by the responses of early level visual neurons tuned to local line segments (e.g., Dickinson, Almeida, Bell, & Badcock, 2010; Fleming, Holtmann-Rice, and Bülthoff, 2011; Dickinson & Badcock, 2013; Bowden et al., 2019; Dickinson, Martin, & Badcock, 2022).

To date, a key obstacle to resolving this debate has been a lack of image-computable models that can make specific, quantitative predictions about which perceived shapes should result from low-level and high-level adaptation processes. Here we directly pit an empirically-derived high-level statistical shape space against a number of local, low-level aftereffect models that are inspired by both neurophysiology (i.e., early stages of visual processing) and previous psychophysical work (e.g., TAE; Bowden et al., 2019).

For the high-level model, we reasoned that the underlying multidimensional ‘shape space’ must encode shape factors (reported previously or still unknown) that are attuned to the statistics of natural shapes. To test this, we probed and modelled high-level shape aftereffects using a recently-developed 22-dimensional statistical shape space (‘*ShapeComp*’), which captures >95% of the variation in a database of ca. 25,000 animal outlines, and which accurately predicts human shape similarity judgments (Morgenstern et al., 2021). Specifically, participants were presented with shapes created using a Generative Adversarial Network (GAN) trained on the animal dataset, and projected into *ShapeComp* space (**Figure 1C**; see **Materials and methods**). This allows creating arrays of novel stimuli that share statistical similarities with the silhouettes of natural forms, within a continuously variable (and approximately perceptually uniform) shape space. For example, starting with an initial coordinate within the ShapeComp space— corresponding to the white, central object in Figure 1C—a 2D array of nearby coordinates yields a set of shapes with continuously varying characteristics. Within this 2D sub-space, we can define a 1D ‘vector trajectory’ of each shape passing from one *adaptor shape* (shapes on the outer rings) through the *mean shape* (white central shape) to the adaptor shape on the opposite side. Lying on the same axis, but on opposite sides of the mean shape, adaptors were approximately equidistant in terms of perceptual similarity from the mean shape, and were thus ‘anti-shapes’ of each other relative to the *mean shape* (cf. ‘anti-faces’; Leopold et al., 2001).

If the *ShapeComp* shape space is a good approximation for human visual shape coding, then adaptation to an adaptor from one end of a vector trajectory should cause biases in subsequent perception of the *test shape* towards the adaptor’s *anti-shape* (**Figure 1AB**). To test this, in each interval of a 2IFC task, observers were peripherally presented (4s) with an adaptor on one side of fixation, and its anti-shape adaptor on the other side of fixation (**Figure 1D** and **Movie S1**). Following adaptation, the first test interval briefly (0.1s) showed two *test shapes* from the vector trajectory, randomly selected to be either identical (i.e., the white *mean shape*) or different. Specifically, the two different test shapes were selected from opposite directions of the mean shape and were equidistant in terms of similarity from the mean shape. Following another round of adaptation, the next test interval showed the remaining *test shapes*, and observers were asked to report which interval contained the pair of shapes that appeared more similar. Having collected responses for a wide range of test and adaptor stimuli, we also compared responses against the predictions of low-level models of adaptation to orientation and position of contour points (see **Figure 2**). To anticipate, we find that the high-level model predicts the perceptual distortions elicited by adaptation substantially better than the low-level aftereffect models.

**Figure 2.**
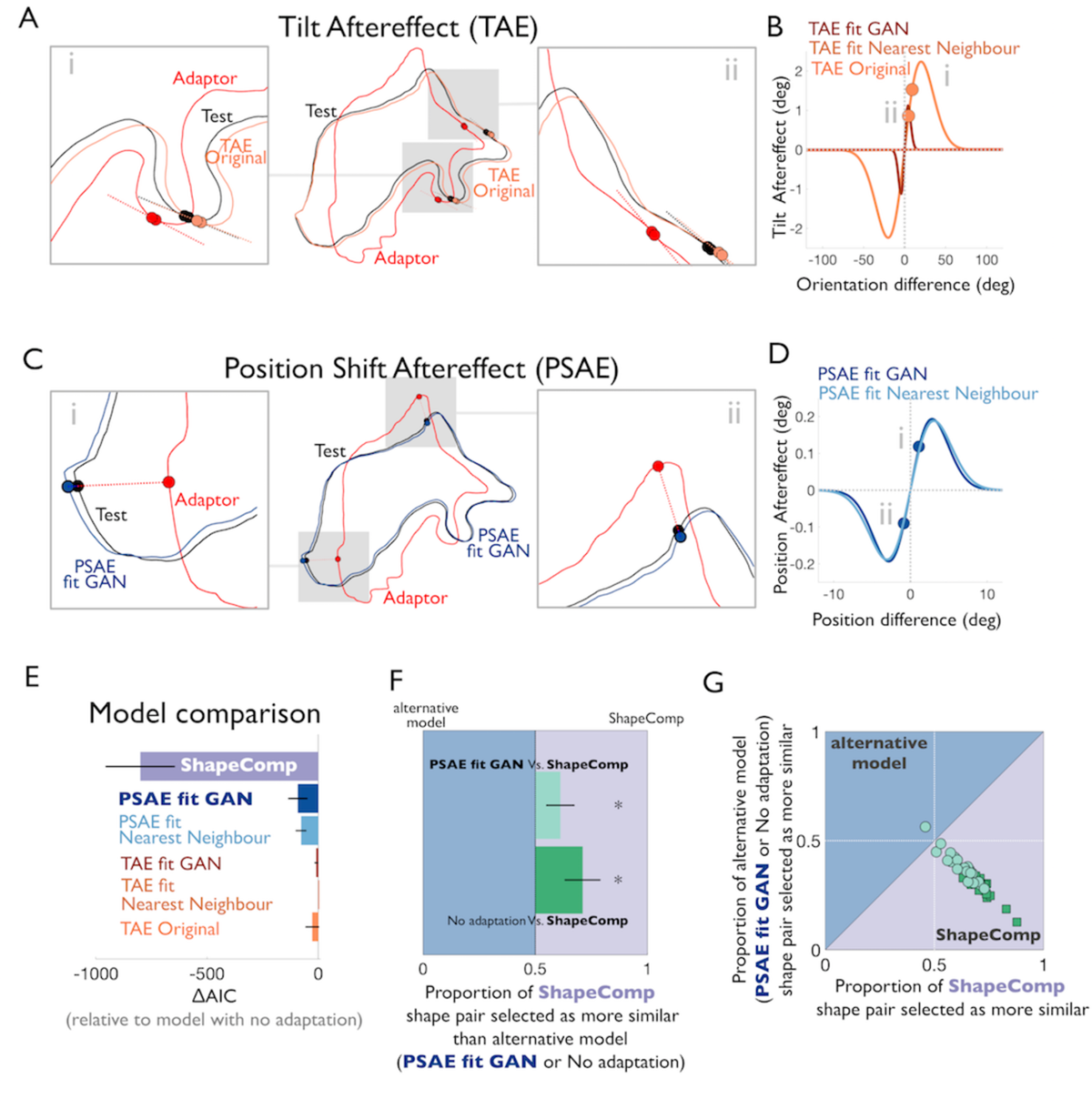
Evaluating local aftereffect models. **(A)** A cartoon depiction of the TAE model where the adaptor (red) perceptually distorts the test shape (black) to produce a predicted aftereffect (orange). The point-to-point correspondence between the adaptor and test shapes is established based on the shape’s generative procedure (see **Figure S3**). Insets highlight examples of line segments inferred by connecting nearby points. After adaptation the corresponding segment of the test shape is perceived tilted away from that segment of the adapting shape. Perceived tilt is computed by using the difference in orientation between the adaptor and test line segments to read off the tilt aftereffect from **(B)** psychophysical data (orange; Bowden et al., 2019). To give local aftereffect models a better chance at predicting the human shape aftereffects, we also implemented versions that were fit to the human responses in Experiment 1 and based on different assumptions about encoding (*TAE fit GAN*, darker red; *TAE fit Nearest Neighbour*; lighter red) (**C**) We also fit two position-shift aftereffect models (*PSAE fit GAN*, darker blue; *PSAE fit Nearest Neighbour*, lighter blue) that predict aftereffects that spatially shift the perceived location of each point on the test shape away from corresponding points on the adaptor. Insets show two examples of position-shifts that are based on computing the difference in spatial position between the adaptor and test points and then reading off the position-shift aftereffect from **(D)** data that were fit to Experiment 1. The new points relative to the test were placed along the same vector that connects the adaptor and test points but away from the adaptor. **(E)** Overall, ShapeComp was much better at predicting the human shape biases from Experiment 1 in terms of a change in the AIC relative to a model with no aftereffects (the *no adaptation* model). With the fitted low-level models in hand, in Experiment 2 we made a fairer comparison and pitted a prediction by ShapeComp against the best local model (the *PSAE fit*) and the *no adaptation* model. With this direct test, we also find that **(F)** observers prefer ShapeComp over the alternative models, (**G**) preferring ShapeComp shape pairs over *PSAE fit* (for 19 out of 20 pairs, light green circles), and over *no adaptation* (for 20 out of 20 shape pairs, dark green squares). Spatial position of data in (**G**) were slightly jittered to reveal overlapping points.

## Results

### Shape aftereffects along vector trajectories in statistical shape space

For Experiment 1, **Figure 1E-G** shows the proportion of times participants selected different test shape pairs as a function of test shape dissimilarity. Negative dissimilarity values refer to test shapes on the opposite side of the mean relative to the adaptor (i.e., anti-shapes, more dissimilar to the adaptor relative to the mean shape), and positive values refer to shapes on the same side of the mean (i.e., more similar to adaptor). If adaption has no effect, observers should rarely, if ever, choose different test shapes over identical mean shapes. Post-adaptation, however, test shapes with positive dissimilarity values were perceived as more similar than identical mean shapes. Thus, adaptation shifted the test shapes towards the mean shapes, and the mean shape towards the adaptor’s anti-shapes (**Figure 1AB**). We found that the test dissimilarity most often selected as looking identical (which we call the ‘shape bias’, peak of black curve in **Figure 1EFG**) was significantly greater than zero across several vector trajectories (0°, 60°, 120° in **Figure 1**), and across independent observers in several different planar slices through shape space (*t*(11) = 7.7, *p*<0.001; **Figure S1**). Thus, adaptation leads to perceptual aftereffects that push shape appearance along specific vector trajectories in the *ShapeComp* space. This provides a first indication that high-level statistical shape coding representations that reflect natural shape variations may provide a plausible account of shape adaptation and thus shape representations more broadly in the human vision.

### Statistical shape space predicts aftereffects better than local models

While the shape aftereffects we observed in Experiment 1 are consistent with adaptation in a high-level multidimensional shape space, it is important to note that they might also be caused by simple low-level processes, such as the encoding of the position (McKee and Westheimer, 1978; Marr, 1982; Geisler, 1984; De Valois and De Valois, 1991) or orientation (Gibson, 1950; Stevens, 1979; Marr, 1982) of local points on the shape’s boundary. This gets to the heart of the long-running debate about whether putative high-level aftereffects might in fact have simple, low-level explanations (Fleming, Holtmann-Rice, and Bülthoff, 2011; Storrs, 2015; Bowden et al., 2019; Dickinson, Martin, and Babcock, 2022). To distinguish between low- and high-level accounts, we first implemented multiple image-computable models of the tilt aftereffect (TAE; Clifford, 2014; Dickinson, Harman, Tan, Almeida, & Badcock, 2012; Gibson & Radner,1937; Mitchell & Muir, 1976; Clifford, Wenderoth, & Spehar, 2000) and position-shift aftereffect (PSAE; Gibson, 1933; Bales and Follansbee, 1935; De Valois and De Valois, 1991; Ramachandran and Anstis, 1990; Whitney, 2002; Maus, Fischer, and Whitney, 2013; Nishida, and Johnston, 1999; McGraw, Walsh, and Barrett, 2004). We then sought the strongest possible low-level model (i.e., the model that best predicts the post-adaptation human shape bias) to compare against a high-level adaptation model based on *ShapeComp* dimensions.

Previously, Bowden et al. (2019) showed that the TAE predicts aftereffects in radial frequency patterns. Bowden et al.’s (2019) model (henceforth ‘*TAE Original’*) predicts perceptual aftereffects by tilting local line segments on a test shape relative to the orientation of nearby local line segments on an adaptor, based on a psychophysically measured TAE function (**Figure 2AB**). To give low-level accounts the strongest possible chance to account for our findings, we also implemented additional versions that were fit to better account for human responses in Experiment 1 and based on different assumptions about encoding (i.e., based on a vector field; **Figure S2**) and point-to-point correspondence (**Figure S3**; i.e., whether based on nearest neighbor, *TAE fit Nearest Neighbour;* lighter red line, **Figure 2B**, or the GAN’s generative procedure, *TAE fit GAN*; darker red line, **Figure 2B**) (See **Materials and methods** local adaptation models for details).

In addition to tilt, adaptation can alter the apparent position of test stimuli (Gibson, 1933; Bales and Follansbee, 1935; De Valois and De Valois, 1991; Ramachandran and Anstis, 1990; Whitney, 2002; Maus, Fischer, and Whitney, 2013; Nishida, and Johnston, 1999; McGraw, Walsh, and Barrett, 2004). Careful inspection of the shape aftereffects we experienced with the stimuli in Experiment 1 led us to suspect that a similar kind of PSAE may have distorted the test shape by shifting the positions of points on its outline away from those on the adaptor. To test this, we therefore also fit a *PSAE fit nearest neighbor* model and a *PSAE fit GAN model* to the human bias data in Experiment 1 (**Figure 2CD**).

We compare the TAE and PSAE low-level aftereffect models to a higher-level one-parameter model from *ShapeComp* statistical shape space. The high-level model’s free parameter was set to predict the aftereffects in Experiment 1 by matching to the mean human shape bias (across 3 vector trajectories in 4 shape spaces from Experiment 1) in *ShapeComp*. We examined which aftereffect model better predicts the human biases in Experiment 1 by computing the Akaike information criterion (AIC) for each model relative to a *no adaptation* model (**Figure 2E**). The *no adaptation* model is a baseline based on the prediction that there are no perceptual aftereffects (i.e., physically identical mean shapes are always seen as most similar post-adaptation). Thus, any model with a negative change in AIC relative to *no adaptation* (ΔAIC) is a better model of the human aftereffects than one that predicts no effect of adaptation.

Indeed, the local aftereffect models are significantly worse at predicting the human bias in Experiment 1 than the one-parameter high-level *ShapeComp* model (**Figure 2E**). However, this comparison is unfair given that the local aftereffect models cause perceptual shifts that do not necessarily match the way the shapes vary along the trajectories of shape space (from Experiment 1; See, for example, **Movie S2**). To provide a more direct comparison of high-level statistical vs. low-level local accounts, in Experiment 2, we directly pitted the best local model, *PSAE fit GAN,* against the statistical shape space, by presenting participants with a choice between the perceived test shapes predicted by each model. Specifically, in each interval of a 2IFC design similar to Experiment 1, adaptors were followed (in a random order) by a prediction of which shape should appear the same in terms of the *ShapeComp* or *PSAE fit GAN* model (See **Figure S4** for example shapes; See **Materials and methods** for details). Observers were again asked to select which of the two intervals contained the more similar pair of shapes, overwhelmingly choosing *ShapeComp* over the *PSAE fit GAN* predictions (*t*(19) = 3.45, *p*<0.001), and also over the no-adaptation model (*t*(19) = 5.19, *p*<0.001; **Figure 2F**). Observers preferred shape pairs from *ShapeComp* over the alternative models for all but one out of a total of 40 shape pairs (**Figure 2G**). Thus, shape aftereffects are more consistent with statistical shape space than simple adaptation to local orientation and/or position.

### Shape aftereffects are attuned to the variation in natural shapes

Lastly, we examined whether these aftereffects are related to a special role for natural geometries in shape encoding. Shape spaces tend to exhibit regularities that reflect the statistics of the datasets from which they are derived. One such regularity is how shapes vary with distance from the origin of the shape space, with central coordinates representing shapes that are more typical of the dataset, and outlying coordinates representing less typical shapes. For example, because *ShapeComp* is derived from animal silhouettes, shapes near the origin tend to be less varied and appear more typically animal-like (compact, smoothly curved geometries with limb-like protrusions, **Figure 3A** *left*) than those further from the origin (bumpy or spiky geometries with strongly elongated aspect ratios, **Figure 3A** *right*). We find that this general tendency (i.e., compact to more elongated shapes) replicates with a different training set of 2000 other animal shapes (‘*replication set’*; **Figure S5A**) but does not necessarily emerge in spaces constructed from artificial shapes (e.g., from shapes based on a random combination of radial frequencies, **Figure S5B**).

**Figure 3.**
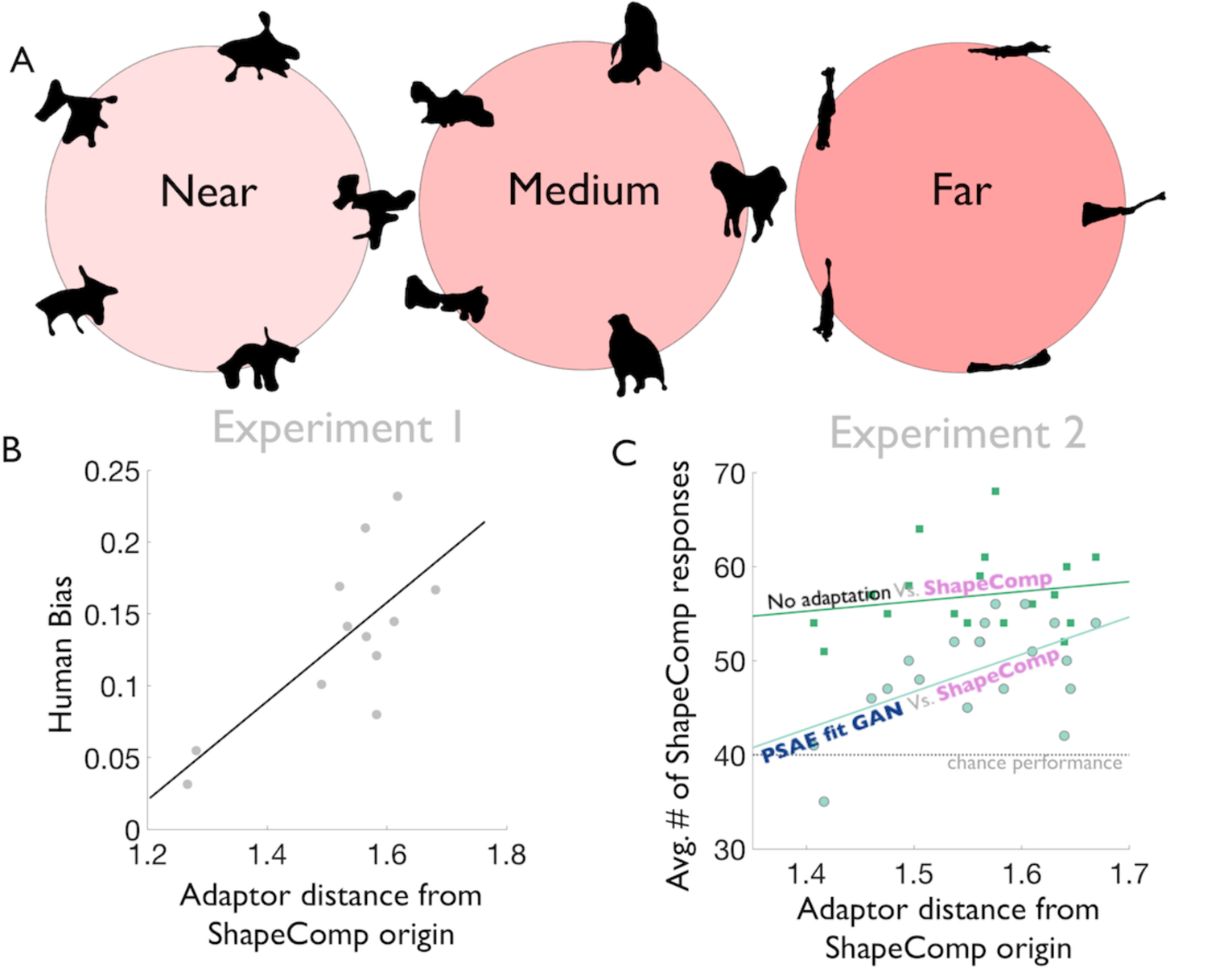
Adaptor distance from origin in statistical shape space predicts strength of aftereffects. (A) GAN shapes relative to the origin of statistical shape space. Example GAN shapes near (left), medium (middle), or far (right) in terms of distance from the ShapeComp origin. Shapes closer to the center are more commonly appearing shape geometries and appear roughly uniform in their aspect ratio. The farther away the shapes are from ShapeComp’s origin, the more novel they are and the less likely the shapes have uniform aspect ratios. If human shape processing is tuned to the commonality of shapes then encoding channels may be more dense and tightly tuned near the centre of the space, where a small geometric change between adaptor and test might equate to non-overlapping encoding channels, and therefore produce little aftereffect; whereas far from origin encoding channels might be broader and more overlapping, so that even large geometric differences will still be encoded by overlapping channels and therefore induce aftereffects on one another. If shape encoding is more precise (i.e., more dense and tightly tuned) at the centre of shape space, then adaptors farther away from the origin cause larger shifts and thus larger perceptual distortions. Indeed, the distance in statistical shape space of adaptor and origin predicts (B) human biases from Experiment 1, and (C) the number of *ShapeComp* preferred responses from Experiment 2, especially in cases when ShapeComp competes with models that are more predictive of shape aftereffects (i.e., when *ShapeComp* is pitted against the *PSAE fit GAN* model). Together, these results suggest that the origin of ShapeComp is a good a priori estimate of a neutral point, and that human shape perception is attuned to the way natural shapes vary.

If human shape coding is attuned to the statistics of natural shapes, then more common shapes that tend to occur nearer to the origin are potentially coded differently than more distant shapes (e.g., more densely or more precisely; Girshick, Landy, and Simoncelli, 2011; Wallis, 2013). If encoding resources are differently allocated depending on the statistics of experienced shapes, we might expect to find different aftereffect tunings associated with different locations in the shape space, as found with color (Eisner and MacLeod, 1981) and face (Webster and MacLin, 1999; Rhodes and Jeffery, 2006) spaces. Moreover, if shape representations are derived from the statistics of natural images, such origin-related effects should be weaker in spaces derived from artificial stimuli than in *ShapeComp* space.

We tested these predictions with our data. Given that the 2D shape slices (in Experiment 1) and shape sets (in Experiment 2) were from randomly selected positions within the shape space, the adaptor distance from the ShapeComp origin varied across our stimuli. This, first, allowed us to re-examine the data of Experiment 1 and 2 to test whether the responses changed as a function of adaptor distance from the origin of *ShapeComp* space. For Experiment 1, we find that adaptors farther away from the origin lead to larger human biases (*r* = 0.74, *p<*0.01; **Figure 3B**). Thus, much like Webster & MacLin (1999) reported that adaptation to distorted faces strongly biases the appearance of a neutral (undistorted) face as compared to adaptation with other neutral faces, adaptation to shapes with more distinctive characteristics tended to yield stronger perceived distortions. For Experiment 2, we compared the proportion of times observers chose *ShapeComp* over competing models, as a function of adaptor distance from the origin (**Figure 2EFG**). As one competing model (the *PSAE fit GAN* model) predicted aftereffects that were more similar to both the ShapeComp prediction and the human bias than the other model (the *no adaptation* model), the comparison between models also allows us to evaluate the strength of the aftereffects (as a function of distance from the origin) under different levels of uncertainty. Under high levels of uncertainty, we find that when comparing *ShapeComp* with *PSAE fit GAN* (**Figure S4**), distance from the origin has an impact (**Figure 3C**, lighter circles; *r* = 0.56, *p<*0.01). Participants selected the *ShapeComp*-predicted shapes more frequently than the *PSAE fit GAN* predictions when the adaptors were distant from the origin, but not when they were close. When the adaptor was near the origin, observers could not easily distinguish between *ShapeComp* and the *PSAE fit GAN* predictions, presumably because the changes between predictions were so small and so were the differences in the aftereffects. When the adaptor was farther from the origin, the predicted differences between the models were still small, but the aftereffects were larger and led to clearer preferences for the ShapeComp predicted distortions. Under lower levels of uncertainty, when competing with a model that does not capture perceptual aftereffects (i.e., *no adaptation* vs. *ShapeComp*), adaptor distance from the origin has less impact in teasing apart pairs of test shapes (**Figure 3C**, darker squares; *r* = 0.19, *p=*0.43); the predicted shape differences between the two models were large, leading to the higher sensitivity and confidence across observers in selecting the ShapeComp predicted distortion regardless of whether the adaptors were near or far from the origin. Thus, in cases of greater uncertainty, when human observers are choosing between models whose predictions are more similar to their own percepts, distance from the origin has a strong impact by shifting response preferences towards empirically-driven statistical shape space. Together these findings suggest different encoding of shapes near the origin of *ShapeComp* space, suggesting a norm derived from the statistics of natural forms.

Second, we tested whether these effects of distance from the origin can be replicated in artificial shape spaces based on radial frequency patterns (**Figure S6A**) or random polygonal geometries (**Figure S6B**). We reasoned that if shape representations are derived from the statistics of natural forms, the relationship between aftereffect strength and distance from the origin should be less pronounced in these arbitrary shape spaces. We therefore re-expressed the data from Experiments 1 and 2 in the coordinates of the artificial shape spaces, and found a substantially weaker dependency of aftereffect strength on distance from the origin. In contrast, the relationship was reproduced in the shape space derived from the abovementioned *replication set*, computed with a different set of 2000 animal shapes than *ShapeComp* (**Figure S6C**). This confirms that results are robust against changes in dataset and, thus, also the noise around the estimate of the origin of shape space. These results highlight an importance of natural shape statistics and, thus, evolution and lifetime learning on the perception and the coding of shape.

Together, these findings suggest that, under higher levels of uncertainty (i.e., when teasing apart models that capture components of the perceptual aftereffects), distance of the adaptor from the origin, which is based on the typicality of the adapting shape within human visual experience, is highly predictive of perceptual aftereffects. These results support the idea of aftereffects as a “re-normalization” or temporary shift in the origin, which allows us to better code differences in shapes based on the statistics of our current environment (Gibson, 1933, 1937; Gibson and Radner, 1937; Helson, 1947, 1964; Rhodes and Jeffery, 2006; Eisner and MacLeod, 1981;Webster and MacLin, 1999; Webster and MacLeod, 2011; Panis, Vangeneugden, Op de Beeck, 2011; Panis, Wagemans, and Op de Beeck, 2011; Kayaert, Op de Beeck, and Wagemans, 2011).

## Discussion

Our findings suggest that shape aftereffects—the distinctive distortions of perceived shape following adaptation—are the result of high-level processes in the visual system that encode shape in a multidimensional feature space, which is derived from the statistics of complex naturalistic forms. Local models based on tilt and position failed to capture the aftereffects we observed as well as the statistical shape space did. Indeed, there are other local features known to be important for shape processing (e.g., information along contour curvature, Feldman and Singh, 2005; local image cues that reveal shape topology, Kunsberg and Zucker, 2021), which may also contribute to shape aftereffects. Statistical shape space, however, encompasses both local and global changes in shape—ShapeComp includes local features such as curvature, and global features such as Fourier descriptors—while also accounting for the way natural shapes vary. Statistical shape space explains phenomena that local process cannot, such as showing many global dimensions (Figure 4 in Morgenstern et al., 2021) previously reported (e.g., *taper*, Suzuki and Cavanagh, 1998; *curvature*, Suzuki, 2001; *aspect ratio*, Regan and Hamstra, 1992; *spikiness*, Haung, 2020; Bao et al., 2020), and explains why some illusory aftereffects are perceived as stronger than others as being a consequence of better shape encoding precision (i.e., more dense and tightly tuned) for more typical shapes which are near the centre of shape space. These results support the role of higher-level shape processes modulating human aftereffects.

An alternate method for testing whether shape aftereffects are high-level is examining whether the aftereffects are non-retinotopic (e.g., Suzuki and Cavanagh, 1998; Bratch et al., 2021). As ShapeComp is a model consisting of both local and global features of shape, we opted to display adaptor and test stimuli in overlapping rather than distinct spatial regions to better drive the encoding process responsible for shape aftereffects. Previously, however, Suzuki and Cavanagh (1998) showed global aftereffects when presenting adaptor and test in retinotopically distinct locations. Whether non-retinotopic shape aftereffects are weaker and by how much is an open question. With statistical shape space, and an understanding that higher-level shape encoding is tuned to shape typicality within human visual experience, we now have additional methods for decoupling lower-level retinopic aftereffects from higher-level global shape factors. Thus, by relying on statistical shape space future work can further expand our understanding of human shape coding in the brain.

Psychophysical adaptation methods have played a key role in characterizing and revealing mechanisms of visual processing (Graham, 1989; Webster, 1996; Lennie and Movshon, 2005), often being referred to as “the psychologist’s microelectrode” (Mollon, 1974; Frisby, 1980; Mather, Verstraten, and Anstis, 1998; Thompson and Burr, 2009; Webster and MacLeod, 2011; Webster, 2015). Many previous studies showed a rough correspondence between psychophysically-measured bandwidths for basic visual properties (e.g., spatial frequency, orientation, and colour) and those in single neurons from the early visual cortex known for coding local information (Lennie and Movshon, 2005). The correspondence has long fueled the debate as to whether shape aftereffects are due to local or global processes (Suzuki, 2005; Storrs, 2015; Bowden et al., 2019). However, another similarity between visual properties that are known to drive adaptation (e.g., orientation, and spatial frequency) is their emergence through statistical summaries of natural patterns, such as efficient coding (Maloney, 1986; Ruderman, Cronin, and Chiao, 1998; Olshausen and Field, 1996; Bell and Sejnowski, 1997) or empirical ranking (Ng et al. 2013; Morgenstern et al., 2014), which also rationalize many aspects of visual perception (Yang and Purves, 2004; Long et al., 2006; Howe and Purves, 2004; Woltach et al., 2008, 2009; Sung et al., 2009). Consistent with this idea, we showed that a statistical shape space—derived from the way animal silhouettes vary, and which predicts human shape similarity (Morgenstern et at., 2021)—also predicts adaptation along vector trajectories in statistical shape space that local processes cannot account for. Such a high-dimensional shape space shows how many shape dimensions from past work (e.g., Regan and Hamstra, 1992; Suzuki and Cavanagh, 1998; Suzuki, 2001; Op de Beeck, Wagemans, and Vogels, 2001; Haung, 2020; Bao et al., 2020) arise in biological shape perception as a means to encode useful regularities in natural image patterns, which are both local and global (i.e., statistical, high-level) in nature. Thus, through evolution and a lifetime of experiences, our brains are highly attuned to the shapes that we regularly see.

## Materials and methods

### Ethics statement

All procedures were approved by the local ethics committee of the Department of Psychology and Sports Sciences of the Justus-Liebig University Giessen (Lokale Ethik-Kommission des Fachbereichs 06, LEK-FB06; application number: 2018-0003) and adhered to the sixth revision of the declaration of Helsinki (2008). All participants provided written informed consent prior to participating.

### Experiment 1: Adaptation along vector trajectories in ShapeComp

#### Participants

*16* participants (11 females; mean age: 25.5 years; range 20–42, 1 author) made shape similarity judgments. All participants were paid 8 Euros per hour, and signed an informed consent approved by the ethics board at Justus-Liebig-University Giessen and in accordance with the Code of Ethics of the World Medical Association (Declaration of Helsinki). All participants reported normal or corrected-to-normal vision.

#### Stimuli

Novel naturalistic shapes were generated with a Generative Adversarial Network (Goodfellow et al., 2020). The GAN was trained in MATLAB using MatConvNet to synthesize novel shapes that the discriminator component of the network could not distinguish from those in a set of >25,000 animal shapes (see Morgenstern et al. (2021) for more details).

We selected vector trajectories through statistical shape space. One possible statistical shape space would be the trained GANs latent vector. However, we opted for using ‘*ShapeComp*’, a statistical shape space computed by summarizing 109 shape descriptors on >25,000 animal shapes, for several reasons. *ShapeComp* is (1) known to be highly predictive of human shape similarity (Morgenstern et al., 2021), and (2) based on features from the computer vision literature whose usefulness in the statistical shape space can be quantified (as opposed to the GAN’s latent space that is based on learning arbitrary features). *ShapeComp* converts a shape’s silhouette into a 22-dimensional vector, where the distance between two shapes in their 22-D representation is predictive of human shape similarity between novel shapes. Morgenstern et al. (2021) provide code to convert a silhouette into the 22-D representation (i.e., Shape-to-ShapeComp). Here we also trained a neural network to convert *ShapeComp* 22-D coordinates to novel GAN shapes (i.e., ShapeComp-to-Shape). Given that there is noise between the mapping of the two solutions, we searched through GAN space and selected vector trajectories that were highly correlated between converting ShapeComp-to-Shape and Shape-to-ShapeComp (*r* = 0.92; 95% CI [0.9 0.94]). Specifically, we used the ShapeComp-to-Shape network and selected four random points in GAN space to serve as different *mean* shapes (i.e., central shape in **Figure 1A**, **S1i**). Each *mean* shape was a common center in a planar 2D slice of statistical shape space that different vector trajectories passed through. For each *mean* shape, we synthesized 3 vector trajectories by selecting sets of adaptors that varied across ShapeComp’s first dimension (0°) or around a combination of the first and second dimensions (60° and 120° counterclockwise from the first dimension) (see **Figure 1A**). Each adaptor set was synthesized on opposite sides of the *mean* shape, with each adaptor positioned about 0.6 ShapeComp units away from the *mean* shape, which observers typically rate as highly similar (Figure 5b from Morgenstern et al., 2021). Test shapes were synthesized to fall between the two adaptors. Finally, we converted the shapes back into ShapeComp (using code from Morgenstern et al., 2021), and selected vector trajectories that were highly correlated between the Shape-to-ShapeComp and ShapeComp-to-Shape solutions.

#### Procedure

All experiments were run with an Eizo ColorEdge CG277 LCD monitor (68 cm display size; 1920 × 1200 resolution) on a Mac Mini 2012 2.3 GHz Intel Core i7 with the Psychophysics toolbox (Kleiner et al., 2007) in MATLAB version 2015a. Observers sat 57 cm from the monitor such that 1 cm on screen subtended 1° visual angle.

Each trial consisted of a 2IFC task, where each interval showed a fixation interval (0.3s) followed by an adaptor interval (4s), followed by another fixation (0.3s) and the test interval (0.1s). The fixation interval showed a fixation dot. The adaptor intervals showed two different adaptor shapes (from either the 0°, 60°, or 120° vector trajectory) positioned to the left and right of the fixation dot. Each adaptor shape was positioned 15.5° from the fixation dot and was spatially jittered in the vertical and horizontal direction every 0.1 seconds around this point by sampling from a random normal distribution (σ =0.6°). On any trial, the adaptor position relative to fixation (whether right or left) was determined randomly. To avoid a weakening of the perceptual aftereffect, different vector trajectories were completed in separate sessions. The test interval showed two mean shapes (physically identical) or two test shapes (physically different) that varied in their similarity (in ShapeComp) along the vector trajectory and symmetrically around the mean shape. The test interval that showed the mean shapes or test shapes was determined randomly. The test interval was accompanied by a beep. The test, mean, and adaptor shapes were roughly 7.25° in spatial extent. The test and mean shapes were peripherally positioned 15.5° from the fixation dot (at the location of the adaptor). Participants were tasked with judging which test interval had the more similar shapes, by pressing 1 for the first interval or 2 for the second interval. The similarity (or ShapeComp distance) between shapes on the test trial was determined by interleaved staircase procedures, which increased test similarity (or decreased dissimilarity) when participants correctly selected the test interval with identical mean shapes.

The duration of each session was roughly 20 minutes. Each participant made shape judgements for a single mean shape, thus completing 3 sessions–one for each vector trajectory. For each of the 4 mean shapes, we recruited 4 different participants.

#### Analysis: Effective human bias

To estimate the effective human bias for each observer in each condition, we found the maximum-likelihood fit of the following curve to the probability of *test* pairs selected as a function of the test dissimilarity:

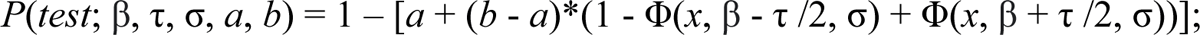

Here Φ(*x*; µ, σ) is the normal cumulative distribution function, *x* is the test dissimilarity, β is the bias (the shape similarity value at which the curve tops out), τ is the width at half-maximum, σ determines the steepness of the curve, and 1 – (*a +* (*b* – *a*)) is the curve’s minimum value. We took β, the value at which test shapes were most likely to be chosen over identical base shapes, to be the effective human bias or maximum strength of perceptual aftereffect for a particular vector trajectory. We constrained the fitted curve at *x* = 0 to pass through 0.5, as this level of test dissimilarity corresponds to a case where the mean shapes are presented in both intervals (which necessarily corresponds to chance performance).

#### Local adaptation models

We posited several models based on two perceptual aftereffects that are known to be spatially localized: the tilt aftereffect (TAE; Clifford, 2014; Dickinson, Harman, Tan, Almeida, & Badcock, 2012; Gibson & Radner,1937; Mitchell & Muir, 1976) and the position-shift aftereffect (PSAE; Gibson, 1933; Bales and Follansbee, 1935; De Valois and De Valois, 1991; Ramachandran and Anstis, 1990; Whitney, 2002; Maus, Fischer, and Whitney, 2013; Nishida, and Johnston, 1999; McGraw, Walsh, and Barrett, 2004) which can be explained by response changes in orientation-tuned neurons (Clifford, Wenderoth, & Spehar, 2000), and spatially-localized neurons, respectively.

#### Tilt After-Effect

The tilt after-effect model is inspired by the work of Bowden et al. (2019). Bowden et al.’s (2019) model predicts shape aftereffects by shifting local points on a test shape relative to an adaptor based on a psychophysically measured tilt-after effect that was fit to a first derivative of a Gaussian:

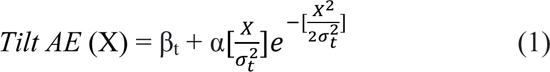

Where X is the orientation difference between the line segments, β_t_ is the Y-intercept (when X =0) and represents a bias in the perception of vertical not induced by the adaptor, α scales the amplitude of the function and σ_t_defines the width of the function. In our implementation, we assumed the point of inflection β_t_ is 0 and the psychophysical parameters averaged across 4 observers from Bowden et al., with α = 74.16°,σ_t_= 20.1°.

Given the human psychophysical function, the TAE model prediction was determined as follows: the local tilt aftereffect for a local orientation difference X (i.e., the angular difference between adaptor (in red) and test or mean shape (in black) at a particular point along the contour; **Figure 2A**) is read-off from a psychophysically measured tilt-aftereffect (**Figure 2B**). This tilt relative to the mean/test is then used to make a local prediction for the position of the perceived shape (e.g., *TAE Original* in **Figure 2A**). In Bowden et al.’s implementation, the predicted shape is determined by an iterative procedure that computes the TAE between the adaptor and mean/test and then lays out the spatial position of the predicted shape one point at a time based on the position of predicted shape’s previous point (Figure 6 from Bowden et al., 2019). We call this model the *TAE Original*.

We also created a version of the model in which the local TAE is treated as a vector field that instantaneously transforms the test/mean shapes into the predicted shape, where the psychophysically measured tilt aftereffect from a local orientation difference is used to infer a vector of given direction and magnitude (blue arrows in **Figure S2**). From these local vectors, we can also extrapolate a global vector field (red arrows in **Figure S2**). One version of the TAE vector field model, *TAE fit GAN*, assumed points were matched based on their default ordering from the GAN–which uses the point ordering to morph smoothly across the latent space (**Figure S3** *left*). In a second version, *TAE fit Nearest Neighbour,* we assumed points were matched based on their nearest neighbor, a strategy that more closely follows the local sensitivities of neurons in early vision (**Figure S3** *right*).

Given that the vector field models are based on a different principle than the *TAE Original* model, we fit parameters for α, and σ_t_ from equation (1) as well as a Gaussian pooling parameter (*SD*), that determines how much TAE to incorporate on the test or mean shape from neighbouring adaptor line segments, such that the *TAE fit GAN* and *TAE fit Nearest Neighbour* best predict the perceptual aftereffects from Experiment 1. Specifically, for each test/mean shape local line segment *L_test_i_*, we gave the most weight to the TAE for the nearest neighbouring adapting local line segment (*TAE fit Nearest Neighbour*) (or the adapting local line segment matched on the default GAN ordering (*TAE fit GAN*)), *L_adapt_i_*, and less weight for TAEs of adapting local line segments (*L_adapt_i+n_* /*L_adapt_i-n_*) that fell on either side of *L_adapt_i_*. The weight was determined with a Gaussian fall-off *N(M,SD)*, where *N* is the normal distribution and M is the point where the adapting and test/mean contour match (i.e., *L_test_i_* = *L_adapt_i_*), and *SD* is the fitted standard deviation. Across all conditions, the models were fit to predict how adaptation leads identical mean shapes to appear more like test shapes towards the adaptor’s anti-shape (by a dissimilarity that matches the human bias, **Figure 1A**). Similar trends were found when fits predicted how different test shapes appear more like the mean (like in **Figure 1B**; See **Figure S7**).

#### Position-Shift After-Effect

In addition to tilt aftereffects, adaptation can bias position percepts (Gibson, 1933; Bales and Follansbee, 1935; De Valois and De Valois, 1991; Ramachandran and Anstis, 1990; Whitney, 2002; Maus, Fischer, and Whitney, 2013; Nishida, and Johnston, 1999; McGraw, Walsh, and Barrett, 2004). Thus, we also fit a *PSAE fit Nearest Neighbor* and a *PSAE fit GAN* to the human bias data in Experiment 1 (**Figure 2CD**), which distorts the test/mean shape to appear to shift away from the adaptor. Like the *TAE fit* and *TAE fit Nearest Neighbour model* above, the fitted PSAE models consisted of 3 parameters: α and σ from equation (1), as well as a Gaussian pooling parameter (*SD*).

### Experiment 2: Evaluating statistical shape space

#### Power analysis

In Experiment 2, we directly pit statistical shape space with (A) *no adaptation*, and (B) the *PSAE fit GAN* model. We used a power analysis to estimate roughly how many participants are needed for each condition. We assumed that if there was no effect between models than observers will be at chance, *H_o_* = 0.5. If observers prefer statistical shape space, then we assumed (1) the estimated effect size in the power analysis should be *less* than the mean of proportion of *test* shapes selected at human bias (or β, threshold location) across the 3 vector trajectories and 4 sets (from Experiment 1), which is 0.61. This is a good estimate of effect size for comparing the *‘statistical shape space’* to the *‘no adaptation’* condition (which is what varied in Experiment 1), but we assumed the effect size to be even smaller for comparisons between ‘*statistical shape space’* and the *PSAE fit GAN* model, *H_1_* = 0.95×0.61 = 0.58. We also assumed (2) the estimated effect size standard error in the power analysis was based on the standard error of the proportion of *test* shapes selected at human bias (or threshold) across the 3 vector trajectories and 4 sets (from Experiment 1), which is 0.083. Assuming a right-tailed *t-test*, we used MATLAB’s *sampsizepwr* function to determine that we need at least 19 participants to reach a power of 0.99 (or alpha level of 0.01). Erring on the side of caution, we decided to collect data from 20 participants.

#### Participants

*20* participants (13 females; mean age: 23.4 years; range 19–30) who were naïve to the purposes and hypotheses of the experiment made shape similarity judgments. All participants were paid 10 Euros per hour, and signed an informed consent approved by the ethics board at Justus-Liebig-University Giessen and adhered to the sixth revision of the declaration of Helsinki (2008). All participants reported normal or corrected-to-normal vision.

#### Stimuli

We synthesized stimuli that pit two models against each other.

### Statistical Shape Space

We searched through ShapeComp for vector trajectories that moved along the spike-like to blob-like dimension known to be an important in macaque (Bao et al., 2020) and human (Huang, 2020) vision. We defined the spike-like to blob-like dimension in ShapeComp using data and stimuli provided by Pinglei Bao (from Bao et al., 2020). We computed ShapeComp for 1224 object shapes from Bao et al. (2020), and then took the spike-like to blob-like vector direction as the direction going from the mean ShapeComp coordinates for the 100 most responsive images from the spike-network (NML, orange dots in Figure 4B from Bao et al., 2020) to the mean ShapeComp coordinates of the 100 most responsive images from the face network (Face, blue dots in Figure 4B from Bao et al., 2020).

With the spike-like to blob-like vector in hand, we then generated shapes with the GAN and searched through statistical shape space for 2 adaptors, 2 tests and a mean shape whose similarity (or distance in ShapeComp units) matched the human bias in Experiment 1, and whose vector trajectory matched the spike-like to blob-like vector. Specifically, we used a genetic algorithm from MATLAB’s global optimization toolbox to search for a shape similarity between (1) the two test shapes and the mean that reflects a distance of 0.25 in ShapeComp space (which is roughly the mean threshold human bias from Experiment 1), (2) the two adaptors that reflects a distance of 1.2 in ShapeComp space, (3) between the two adaptors and the mean shape that reflects a distance of 0.6 units in ShapeComp space, and (4) between the two adaptors and their nearest test shape that reflects a distance of 0.35 units in ShapeComp space. In addition, we also minimized the error between the correlations of spike-like to blob-like vector and the vector direction between (A) the adaptors, (B) the adaptors and the test shapes, and (C) the adaptors and the mean shape. Using this procedure, we created 20 different sets of adaptor, test, and mean shapes, ensuring that the final correlation between the vectors in (A), (B), and (C) and the spike-like to blob-like vector is at least *r =*0.79 (r^-^ = 0.84, σ_t_=0.01). In one session, we pitted the predicted test shapes against the mean shapes–*no adaptation* vs. *ShapeComp*. In another session, we pitted the predicted test shapes against the *PSAE Fit* model (whose synthesis is described below)–*PSAE fit GAN* Vs.

### ShapeComp

#### Position-Shift After-Effect Mode

For each adaptor in the 20 sets synthesized in the last section, we used the *PSAE fit GAN* model from **Figure 2** to find a test stimulus that produces an aftereffect that makes the test appear like the mean shape. Specifically, we used a genetic algorithm to find the least squares error between corresponding points of the *PSAE fit GAN* prediction (of the adaptor on a test) and the mean shape.

#### Procedure

Each trial consisted of a 2IFC sequence that repeated 3 times per trial to increase the chance that observers saw the briefly presented test. Before each 2IFC sequence, the repetition number (1, 2, or 3) was presented centrally at fixation (0.75s). Afterwards, each interval of the 2IFC sequence showed an initial fixation interval (0.3s) that preceded an adaptor interval (2s), followed by another fixation (0.3s) and the test interval (0.1s). The fixation interval showed a centrally presented fixation dot. The adaptor intervals showed two different adaptor shapes (along the spike-like to blob-like vector trajectory; described in *Stimuli: Statistical Shape Space*) positioned to the left and right of the fixation dot. The location of the adaptor relative to fixation (i.e., whether right or left) was kept constant over a single trial, but was determined randomly across trials. Like Experiment 1, the mean position of each adaptor shape was 15.5° from the fixation dot and was spatially jittered in the vertical and horizontal direction every 0.1 seconds around this position by sampling from a random normal distribution (σ =0.6°). The test interval showed the test shapes, which were the predictions from *statistical shape space* or one of the other models (i.e., *no adaptation* or *PSAE fit GAN*). Within the 2IFC sequence, the test interval that showed test shapes from either *statistical shape space* or the competing model was determined randomly. The test interval was accompanied by a beep. All shapes were roughly 7.25° in spatial extent. The test interval shapes were peripherally positioned 15.5° from the fixation dot (at the mean location that the adaptor jittered around). Participants were asked to judge which test interval had the more similar shapes, by pressing 1 for the first interval or 2 for the second interval.

To get accustomed to the trial sequence, before the first session, observers ran through 5 example trials (using stimuli that were not repeated in the main experiments). In one 25 minute session, *statistical shape space* (or *ShapeComp*) was pitted against *no adaptation:* the test interval showed two mean shapes (physically identical) or two test shapes (physically different) that varied in their similarity (in ShapeComp) along the spike-like to blob-like vector trajectory and symmetrically around the mean shape and whose distance from the adaptor along the vector trajectory was based on the human bias from Experiment 1 (as described above in *Stimuli: Statistical Shape Space)*. In another 25-minute session, ‘*statistical shape space’* (or *ShapeComp*) was pitted against the *PSAE fit GAN* model: the test interval showed the predicted test shape from *PSAE fit GAN* model that, post-adaptor, causes the test shapes appear to like the mean shapes or two test shapes that varied in their similarity (in ShapeComp) along the spike-like to blob-like vector trajectory (as described in *Stimuli: Statistical Shape Space*). The sessions (i.e., whether *statistical shape space* was pitted against *no adaptation* or the *PSAE fit GAN* model) were determined in a random sequence. Each session consisted of 80 trials; 20 sequences of stimuli that varied along the spike-like to blob-like dimension that were repeated 4 times each.

### Analyzing the most relevant dimensions

Given that the vector trajectories of the 20 stimulus sets were not identical but each honed in on the spike-like to blob-like dimension, we compute adaptor distances from ShapeComp’s origin based on a subset of the dimensions assuming that an observer’s adaptation processes take account the most apparent shape changes. We define the most relevant dimensions of the unit vector passing through the spike-like to blob-like trajectory as scalar components with the largest magnitude, which also cause the greatest change along the trajectory. We computed these dimensions by sorting the scalar components by their absolute magnitude and then selecting those components whose sum of squares is nearest to 95% of the length of the unit vector. This yielded 4 out of 22 ShapeComp dimensions, which is an efficient shape description of the spike-like to blob-like trajectory. We evaluated the distance of each adaptor to the origin (in **Figure 3C**) based on these 4 dimensions.

### Natural and non-natural shape spaces

#### Stimuli

We compare statistical shape space based on ShapeComp with two artificial shape spaces that share commonalities with natural shapes, radial frequencies and polygonal edges. Radial frequencies share the local smoothness and curvature of natural shapes. Polygonal edges, on the other hand, are jagged, but their environments are constructed to capture how typical natural shapes show roughly uniform aspect ratios.

#### Radial frequency (RF) patterns

Radial frequency (RF) patterns are a special class of closed contours defined by modulation of a circle’s radius. The radius at any point on the contour can be described as follows:

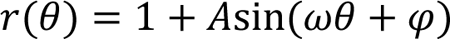

where *r* and θ represent each point on the contour in polar coordinates, *A* is the amplitude of deformation, and ω is the number of cycles of modulation in 360**°**. RF patterns are commonly used to examine shape coding (e.g., Habak, Wilkinson, and Wilson, 2006; Loffler, 2008; Bell and Badcock, 2009; Loffler, 2015) and adaptation (e.g., Anderson et al., 2007; Bell, Dickinson, & Badcock, 2008; Bowden et al., 2019).

We synthesized environments that combined 10 radial frequency patterns, where the parameters *A,* θ, and ω where sampled from varying random distributions (example shapes in **Figure S5A**). RF patterns were constrained to be without self-intersections.

#### Polygonal edge (PE) patterns

We also synthesized straight edge polygons that consisted of 8 to 10 vertices (determined randomly) whose radial distance from the unit circle varied uniformly from [0,1], and whose angle was chosen to fall around a unique bin from [0° to 324°, with a step size of 36°]. Shapes were constrained to be without self-intersections. Like statistical shape space based on natural shapes, the polygonal edge patterns created in this way render environments whose most common shapes are symmetric, and most uncommon shapes are extended (**Figure S5B**).

#### Natural animal shapes

Given that the results in **Figure 3BC** are based on a neural network trained to predict ShapeComp coordinates with a small amount of error, we chose to validate the results. Thus, we also took random samples of animal shapes from an animal shape dataset used in Morgenstern et al. (2021). This is the *replication set*.

#### Shape spaces

For each shape type (RF, PE, or natural animal), we rendered 4 environments with 2000 shapes. We computed shape space following the methods described in ‘Real-world shape analysis’ from Morgenstern et al. (2021): We computed Euclidean distance between each pair of shapes in the dataset each shape with 109 shape descriptors for the computer vision literature, and then we used multidimensional scaling to find orthogonal set of shape dimensions that captures unique variance across the shape set. **Figure S5** shows examples of typical shapes (near the origin of shape space) and atypical shapes (farther away from the origin).

#### Comparing perceptual aftereffects across shape spaces

We examined how well the structure of these shape spaces account for human perceptual aftereffects from Experiment 1 and 2. To do this, we had to determine the coordinates of adaptor shapes in the new shape spaces (see ‘Estimating coordinates for new shapes in pre-existing shape spaces’ in Morgenstern et al., (2021)). With the coordinates in hand, we compared how well distance of the adaptors from the origin predicted the strength of perceptual aftereffects (**Figure S6**). Specifically, for Experiment 2, given that adaptors varied from trial-to-trial within a session, we computed the distance across the most relevant dimensions along the spike-like to blob-like dimension for each shape space (see *Analyzing the most relevant dimensions*). We did not do this for Experiment 1 given that, with the same adaptors seen throughout a session, there is less noise in the adaptation processes.

## Acknowledgements

We thank Saskia Honnefeller and Annika Zentel for their help setting up the experiments and running initial pilot studies.

## Funding

This work was funded by the Deutsche Forschungsgemeinschaft (DFG, German Research Foundation)–project number 222641018–SFB/TRR 135 TP C1, by the European Research Council (ERC) Consolidator Award ‘SHAPE’–project number ERC-CoG-2015-682859, by Research Cluster “The Adaptive Mind” funded by the Hessian Ministry for Higher Education, Research, Science and the Arts, and a Marsden Fast Start grant from the Royal Society of New Zealand awarded to KS (project number MFP-UOA2109).

---

**Figure S1.**
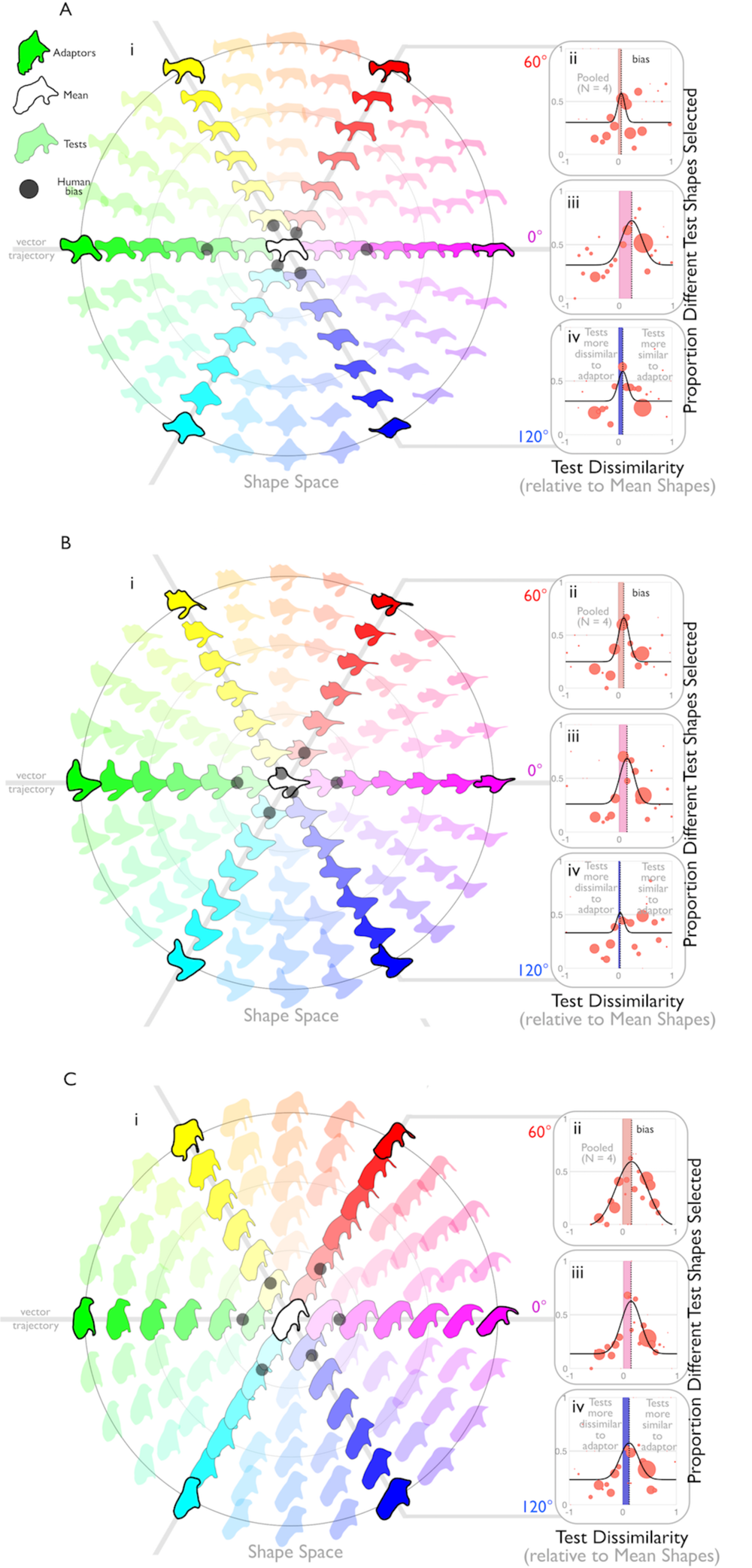
Experiment 1, other shape sets: Adaptation along vector trajectories in statistical shape space yield perceptual aftereffects. A (A) second, (B) third, and (C) and forth set of GAN shapes projected onto a slice of statistical shape space (‘*ShapeComp*’). (i) To highlight shape similarities, shapes are represented in a continuous HSV colour space, where hue changes reflect the angular component of the slice, and saturation reflect similarity from the central mean shape. In separate sessions, our adaptation experiment selected stimuli that passed through single vector trajectories in shape space (at 0°, 60°, or 120°), with the adaptors on the outer ring, and test shapes falling between them. In the 0°, 60°, and 120° (ii,iii,iv) conditions, we find that after the adaptor interval there exist a range of test shapes that observers prefer as more similar than identical mean shapes. The black curve is a maximum-likelihood fit of a double-sigmoid function through the red data points (see Materials and methods for details). We call the test dissimilarity that leads to the highest probability of selecting different test shapes over identical mean shapes (i.e., peak of the black curve) as the ‘shape bias’, which is also shown for each vector trajectory as achromatic dots in (i).

**Figure S2.**
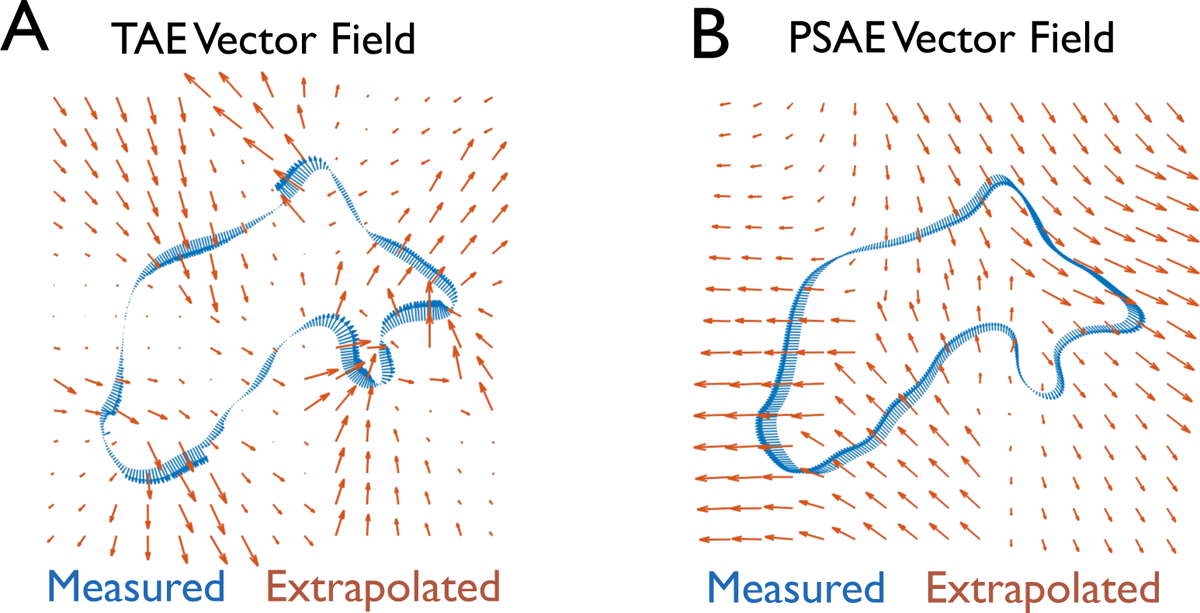
The vector field models. Examples of (A) TAE and PSAE (B) vector fields when correspondence matching is based on the GAN generative procedure. The measured vector fields are in blue, and extrapolated vector fields in red.

**Figure S3.**
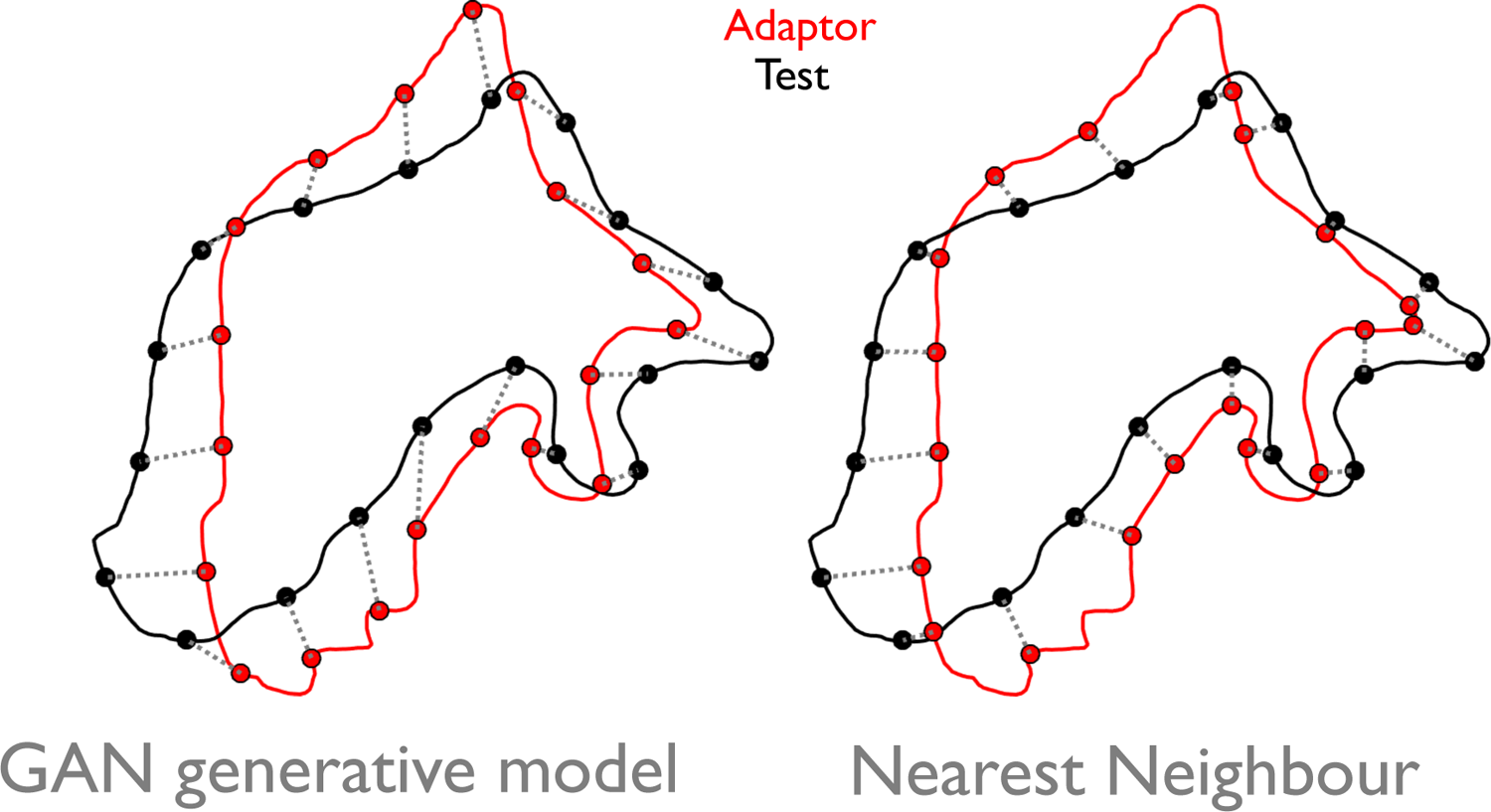
Different point-to-point correspondence matching in models. Point-to-point correspondence between adaptor (red) and test (black) shapes based on the GANs generative model (left) and based on finding the nearest neighbour point on the adaptor for a given local point on the test shape (right). The generative matching strategy is in a sense smarter as it matches the points based on their corresponding parts (e.g., the top hump on the adaptor with the top hump of the test). The nearest neighbour rule is more consistent with the workings of local processes in early vision.

**Figure S4.**
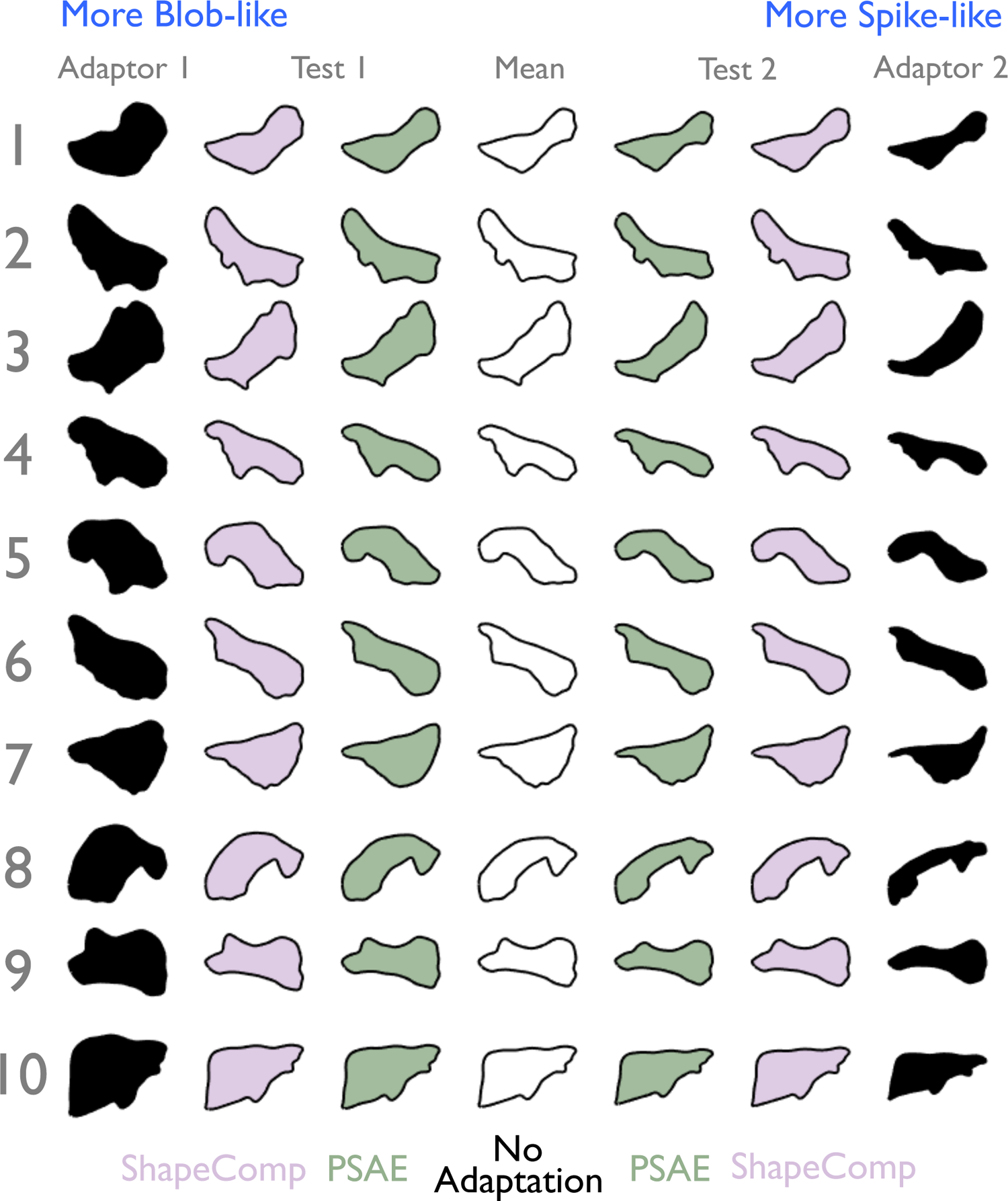
Example stimuli from Experiment 2. Adaptor, test, and mean shapes for 10 out of 20 sets of stimuli that varied along the spike-like to blob-like dimension in statistical shape space. In one session, the test stimuli pitted the *PSAE fit GAN* model (green) against *ShapeComp* (light purple). In another session, the test stimuli pitted the *no adaptation* model (white) against *ShapeComp*. Adaptors are filled in with black. Test 1 followed Adaptor 1, and Test 2 followed Adaptor 2. In the *no adaptation* condition, identical mean shapes followed both adaptors. See **Materials and methods** for details.

**Figure S5.**
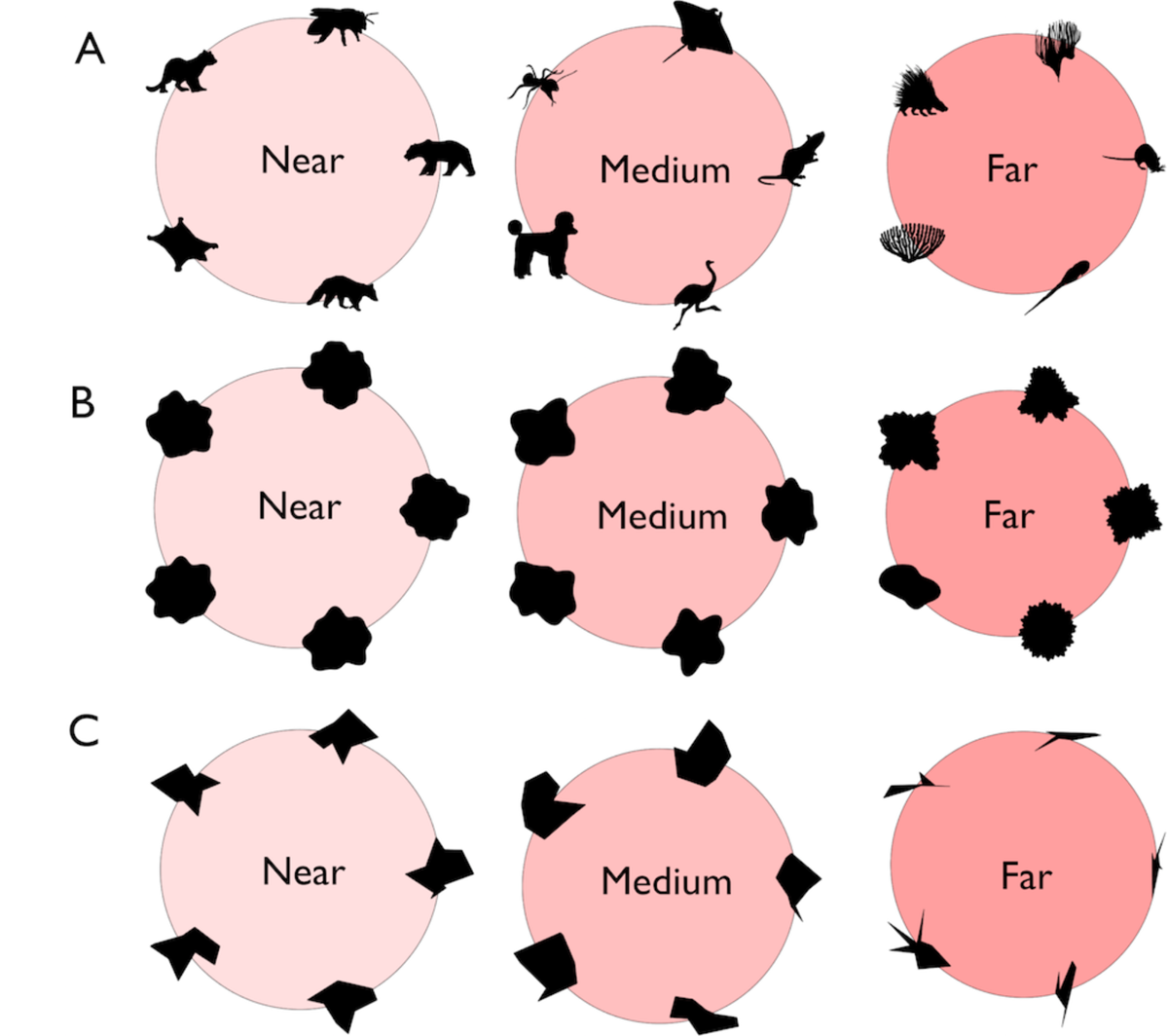
Shape relative to the origin of their shape space. Example (**A**) animal shapes from the ‘*replication set’* (**B**) radial frequency (RF) and (**C**) polygon edge (PE) shapes near (left), medium (middle), or far (right) in terms of distance from the origin of their respective shape spaces. Shapes closer to the center are more commonly appearing shape geometries from that space, and farther away shapes are less likely. The RF shapes (in **B**) appear to consist of a mid-range of radial frequencies near the origin in comparison to shapes that are medium or far distances from their origin, which begin to show more coarse or fine radial frequencies. While the RF shapes share the local smoothness and gradually varying curvature of natural shapes, non-typical RF shapes are not as asymmetric as non-typical natural animal shapes. On the other hand, PE shapes are much more jagged in comparison to natural shapes (in **A**), but like natural shapes, typical PE shapes (in **C**) appear as roughly uniform in their aspect ratio in comparison to non-typical shapes that become more spiky and elongated.

**Figure S6.**
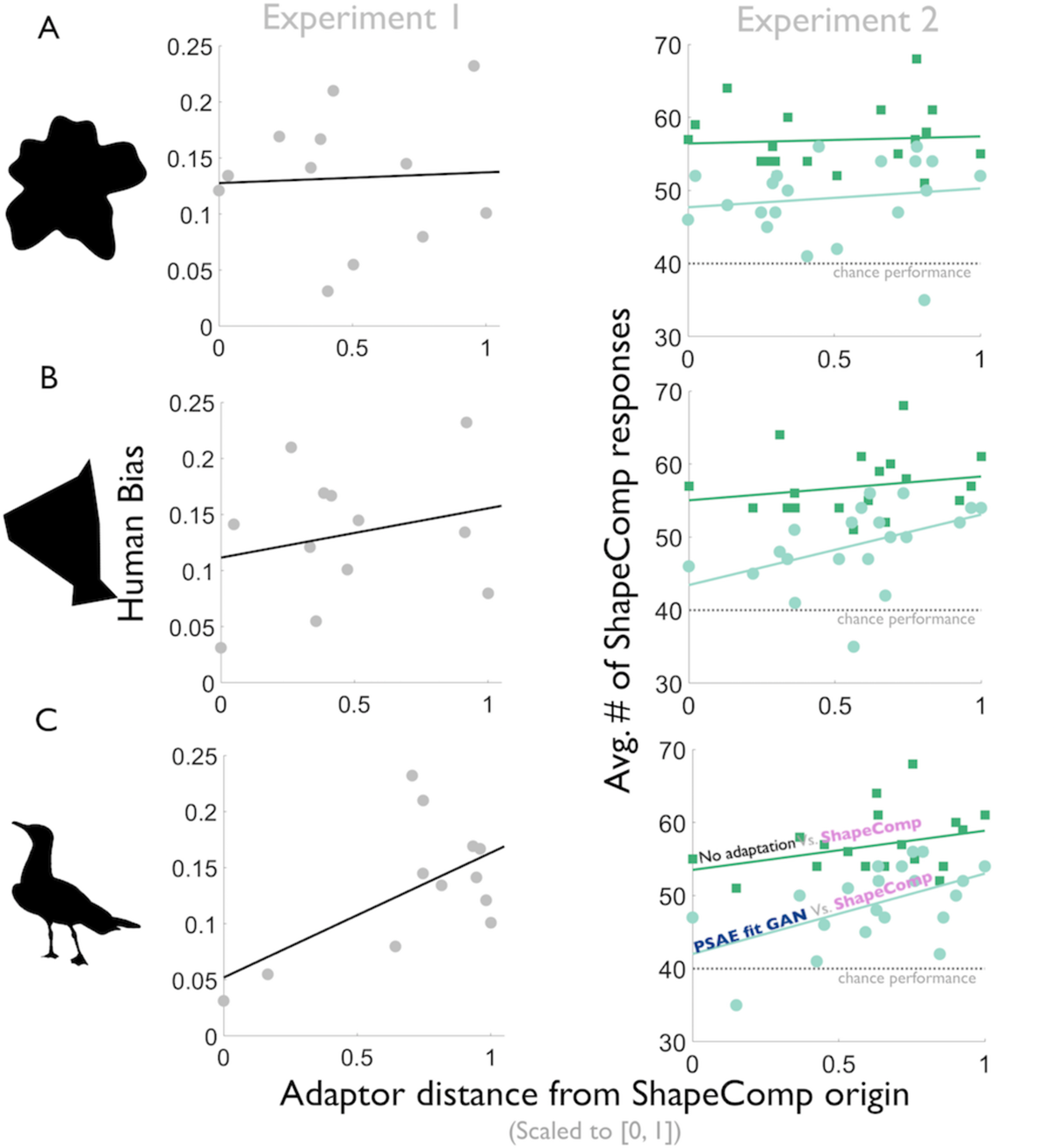
The structure of natural shape spaces predicts aftereffects better than of non-natural shape spaces. (**A**) The distance in statistical shape space for an artificial shape space based on a combination of radial frequency patterns does not predict the human biases in Experiment 1 (*r* = 0.05, *p* = 0.87) and Experiment 2 (*PSAE fit GAN* vs. *ShapeComp*: *r* = 0.14, *p* = 0.55; *no adaptation* vs. *ShapeComp*: *r* = 0.07, *p* = 0.78), suggesting that natural shape statistics that go beyond locally smooth and continuous contours plays an important role in shape coding. Presumably, the roughly uniform aspect ratio of typical shapes, and the large asymmetry of shapes farther away from the origin of shape space is an important factor (Figure 3A). (**B**) Consistent with this idea, the distance between adaptor and origin in shape spaces from random polygonal edges (which also show atypical shapes as more asymmetric, see **Figure S5B**) is better predictive of the human aftereffects in both Experiment 1 (*r* = 0.24, *p* = 0.45) and Experiment 2 (*PSAE fit GAN* vs. *ShapeComp*: *r* = 0.45, *p* = 0.045; *no adaptation* vs. *ShapeComp*: *r* = 0.19, *p* = 0.41). (**C**) Shape aftereffects in natural shape spaces based on animal shapes are most predictive of the human aftereffects in both Experiment 1 (*r* = 0.60, *p* = 0.04) and Experiment 2 (*PSAE fit GAN* vs. *ShapeComp*: *r* = 0.52, *p* = 0.02; *no adaptation* vs. *ShapeComp*: *r* = 0.32, *p* = 0.16), replicating the trends from Figure 3BC. For each environment (radial frequency, polygonal animal, or animal), average adaptor distance from shape space origin is computed from 4 separate shape spaces computed with 2000 randomly generated (or randomly selected) shapes. Together, these results suggest that shape spaces based on artificial shapes do not necessarily capture the structure and variation of natural shapes, and that human shape perception is better attuned to natural shapes spaces.

**Figure S7.**
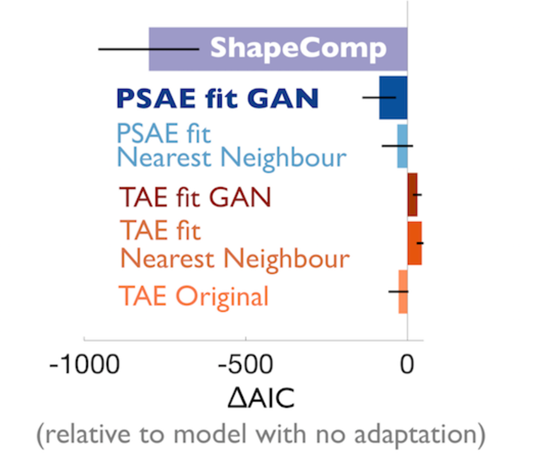
Local model evaluation. In the main manuscript we evaluate models that predict mean shapes were shifted along the vector trajectories in the opposite direction of the adaptor (as in Figure 1A). We find a similar trend when we evaluate models that predict post-adaptation test shapes look more like the mean shape (as in Figure 1B). The *PSAE fit GAN* (based on Figure 1B) is still the best local model, and produces predicted aftereffects for Experiment 2 that are nearly identical to the *PSAE fit GAN* (based on Figure 1A).

## Supplemental Materials

**Movie S1.** Example trial from Experiment 1.

**Movie S2.** Changing the low-level model parameters (e.g., PSAE fit GAN) cause aftereffect predictions that are distinct and very different than ShapeComp’s predicted aftereffect. Adaptor (green), test (black), and predicted shape (filled in with gray).

## Notes

### Competing Interest Statement

The authors have declared no competing interest.

